# KNL1 and NDC80 represent new universal markers for the detection of functional centromeres in plants

**DOI:** 10.1101/2023.12.21.572763

**Authors:** Ludmila Oliveira, Pavel Neumann, Yennifer Mata-Sucre, Yi-Tzu Kuo, André Marques, Veit Schubert, Jiří Macas

## Abstract

Centromere is the chromosomal site of kinetochore assembly and microtubule attachment for chromosome segregation. Given its importance, markers that allow specific labeling of centromeric chromatin throughout the cell cycle and across all chromosome types are sought for facilitating various centromere studies. Antibodies against the N-terminal region of CENH3 are commonly used for this purpose, since CENH3 is the near-universal marker of functional centromeres. However, because the N-terminal region of CENH3 is highly variable among plant species, antibodies directed against this region usually function only in a small group of closely related species. As a more versatile alternative, we present here antibodies targeted to the conserved domains of two outer kinetochore proteins, KNL1 and NDC80. Sequence comparison of these domains across more than 350 plant species revealed a high degree of conservation, particularly within a six amino acid motif, FFGPVS in KNL1, suggesting that both antibodies would function in a wide range of plant species. This assumption was confirmed by immunolabeling experiments in angiosperm (monocot and dicot) and gymnosperm species, including those with mono-, holo-, and meta-polycentric chromosomes. In addition to centromere labeling on condensed chromosomes during cell division, both antibodies detected the corresponding regions in the interphase nuclei of most species tested. These results demonstrated that KNL1 and NDC80 are better suited for immunolabeling centromeres than CENH3, because antibodies against these proteins offer incomparably greater versatility across different plant species which is particularly convenient for studying the organization and function of the centromere in non-model species.

## Introduction

Centromere is a site on chromosomes that mediates their attachment to spindle microtubules, thus playing a crucial role in chromosome segregation during cell division. Cytogenetically, the centromere can often be recognized as a primary constriction on mitotic chromosomes. However, the applicability of chromosome morphology to the determination of centromere location is limited, because genuine functional centromere domains make up only a fraction of the chromatin in primary constrictions. Moreover, primary constrictions are visible only on condensed chromosomes, are difficult to discern on small chromosomes, and are missing on holocentric chromosomes, which have centromere domains distributed along the entire chromosome length (Schubert et al. 2020). Investigation of the size of centromeres as well as their chromosomal and nuclear organization and DNA sequence composition requires the accurate identification of centromere domains during the entire cell cycle, which is only possible using centromere-specific molecular markers.

Centromeric DNA sequences vary considerably among species and, in some cases, among individual chromosomes of the same species or between centromere domains on the same chromosome (Houben and Schubert 2003; Neumann et al. 2012; Oliveira and Torres 2018; Ávila Robledillo et al. 2020). Furthermore, centromeric DNA sequences can also be found in non-centromere locations. Therefore, the localization of centromeres, based on their nucleotide sequence, is limited to species with known centromere DNA composition and those containing centromere-specific DNA sequences.

In contrast to the nucleotide sequence, the protein sequence composition of centromeres is highly conserved and comprises mainly kinetochore and kinetochore-associated proteins (Schalch and Steiner 2017). Kinetochore is a complex multiprotein structure that forms specifically on centromere domains and connects centromeric chromatin with spindle microtubules. The foundational kinetochore protein is CENH3, a centromere-specific variant of histone H3 that replaces the canonical H3 in centromeric nucleosomes (McKinley and Cheeseman 2016). The amino acid sequences of the histone-fold domains of CENH3 and H3 are similar at the C-terminus but differ at the N-terminus (Malik and Henikoff 2001; Talbert et al. 2002; Jiang et al. 2003). Antibodies against N-terminal tails of CENH3 histones have become the most widely applied means to detect functional centromere domains, not only in plants but also in fungi and animals. However, because the N-terminus of CENH3 is highly variable among species, the antibodies directed against this region either recognize CENH3 only in the species in which they were developed or, at most, in closely related species. Consequently, anti-CENH3 antibodies need to be developed repeatedly for each group of closely related species, which is time-consuming and expensive. Although antibodies against other kinetochore proteins such as CENPC (Dawe et al. 1999; Hoopen et al. 2000), NDC80 (Du and Dawe 2007), and KNL1 (Su et al. 2021) have been developed for some plant species, none of them have proven to be more universal plant centromere-specific markers than CENH3.

A commercial antibody against histone H2A phosphorylated at threonine 120 (H2AT120ph) has been the most versatile antibody, to date, for labeling (peri)centromeric regions (Demidov et al. 2014). Although H2AT120ph is present in centromeres, its localization relative to CENH3-containing domains differs among species and between mitosis and meiosis in some species (Cabral et al. 2014). Thus, although H2AT120ph is a useful centromere marker for species for which no other centromere-specific antibodies are available, it cannot fully replace CENH3, which defines functional centromere domains more precisely (Neumann et al. 2016).

In our previous study, which focused on the composition of kinetochore proteins in the monocentric and holocentric *Cuscuta* species (Eudicotyledons, Convolvulaceae), we developed rabbit polyclonal antibodies against the structural kinetochore proteins KNL1 and NDC80 (Neumann et al. 2023). Peptide immunogens used for developing antibodies against these two kinetochore proteins were designed based on domains that were conserved between monocentric and holocentric *Cuscuta* species and that showed a high level of similarity to homologous proteins in *Ipomoea* spp. (Convolvulaceae) and the evolutionarily distant *Arabidopsis thaliana* (Brassicaceae). In situ immunodetection experiments revealed that the developed antibodies functioned not only in *Cuscuta* species but also in the evolutionarily distant *Rhynchospora pubera* (Monocotyledons, Cyperaceae), which was included in the study as a holocentric control species. These results indicated that the developed antibodies would likely detect target proteins in a wide range of angiosperms (flowering plants); however, the extent of their reactivity with KNL1 and NDC80 proteins in other plant species remained unexplored.

In this study, we aimed to explore the range of the reactivity of anti-KNL1 and - NDC80 antibodies by determining the sequence divergence of domains selected for peptide immunogens in *Cuscuta* spp. and by performing in situ immunodetection of the two proteins in species that showed different degree of sequence similarity of these domains and represented plant lineages with different evolutionary distances from *Cuscuta*. Our primary focus was on angiosperms, because they were most likely to have sufficient sequence similarity to peptide immunogens used for developing the antibodies in *Cuscuta* species, but we also analyzed sequences from gymnosperms and several non-seed plant species as outgroups. Our results suggested that anti-KNL1 and -NDC80 antibodies are likely to function in highly diverse plant species. Additionally, the anti-KNL1 antibody recognized a motif that is fully conserved in the majority of seed plants, indicating that this antibody is likely to be highly versatile.

## Results

### KNL1 and NDC80 immunogen domains show high similarity to homologous sequences from a wide range of seed plants

To assess the sequence variability of domains used as peptide immunogens for developing anti-KNL1 and -NDC80 antibodies in our previous study (Neumann et al. 2023), we performed a large-scale screening of homologous proteins in seed plants. Most protein sequences were identified using iterative blastp searches against the protein sequence database in GenBank. For some species from poorly represented lineages with an available whole-genome sequence assembly but inadequate or unavailable gene/protein predictions, we performed tblastn searches to find at least the domains corresponding to the peptide immunogens. In total, we gathered sequence data from 383 species, including 355 angiosperms (from 84 families), 19 gymnosperms (five families), and eight non-seed plants (six families) (Fig. S1, Tables S1 and S2). In 345 of these species (90.1%), we found both KNL1 and NDC80 genes.

Comparison of the KNL1 peptide immunogen with KNL1 sequences from seed plants revealed 40–90% sequence identity (average 76.1%) in angiosperms and 30–50% sequence identity (average 42.7%) in gymnosperms (Table S1). A conserved amino-acid sequence motif (FFGPVS) was found in 276 species of angiosperms (83%) and all 19 species of gymnosperms included in this study (Fig. 1, Fig. S1, and Table S1). Considering that peptide immunogens usually elicit antibodies that bind linear epitopes of 4–12 amino acid residues in length (Buus et al. 2012) and that the FFGPVS motif was the only stretch of ≥4 amino acid residues that was identical between the immunogen sequence and KNL1 from *Rhynchospora pubera*, the good performance of anti-KNL1 antibody observed previously in *R. pubera* (Neumann et al. 2023) suggested that the motif is a possible epitope. If confirmed, this would predict a high versatility of the anti-KNL1 antibody for centromere labeling in seed plants. On the other hand, non-seed plant species showed only 5–25% identity to the domain and lacked the FFGPVS motif, suggesting that their KNL1 proteins are unlikely to be recognized by the antibody (Fig. 1 and Table S1).

**Fig. 1.**
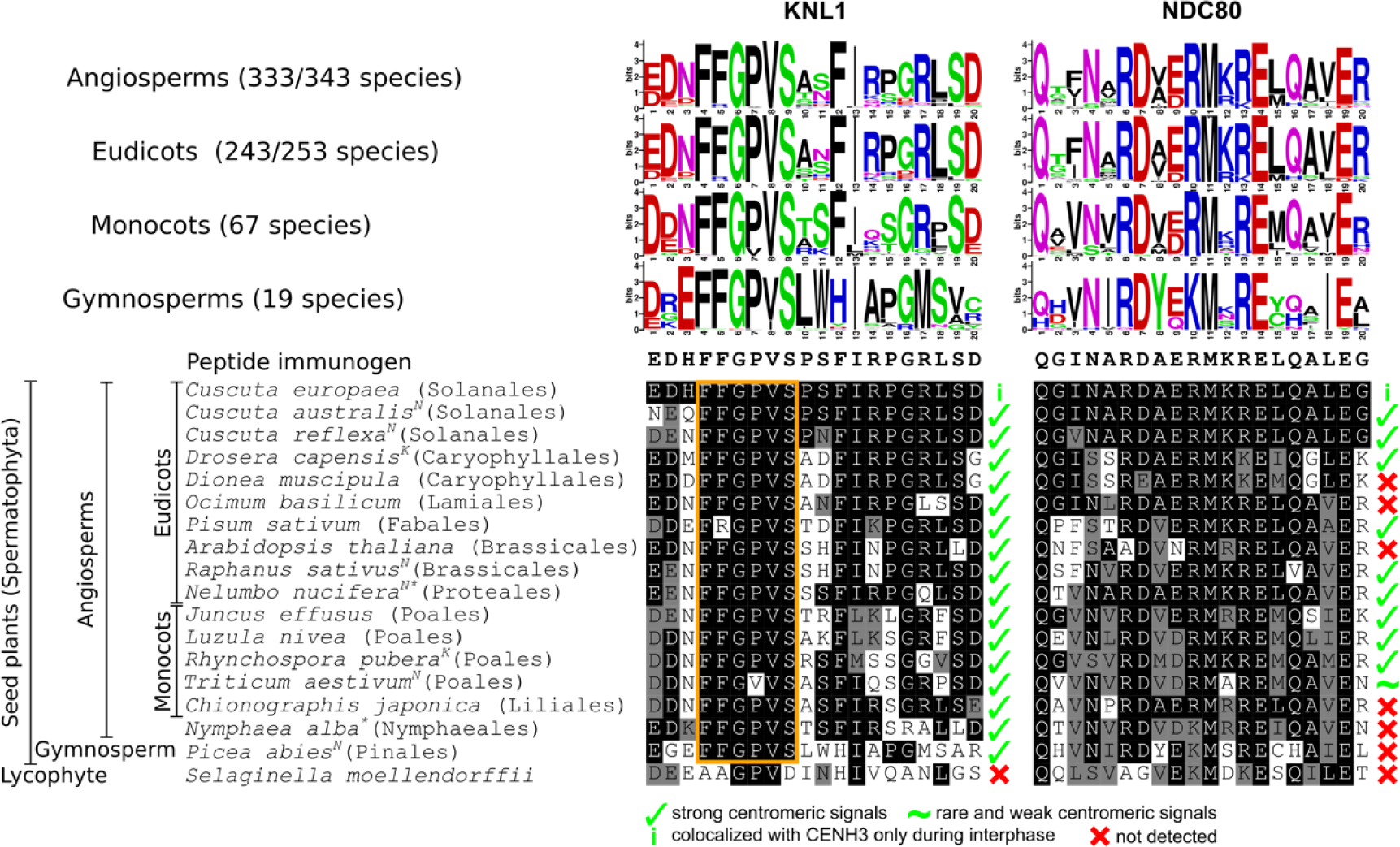
Comparison of the amino acid sequence of the domains used as immunogen for developing anti-KNL1 and -NDC80 antibodies with that of the corresponding domain in other seed plants. Superscript K and N indicate the presence of variants of *KNL1* and *NDC80* genes, respectively. Asterisks indicate the basal eudicot (*Nelumbo nucifera*) and basal angiosperm (*Nymphaea alba*) species. Black and gray box-shading indicates amino acid residues that are identical and similar, respectively, to the peptide immunogens used for developing the antibodies against KNL1 and NDC80. Orange box highlights a conserved motif within the KNL1 domain.

The immunogen sequence of NDC80 shared 40–100% identity (average 69.5%) with angiosperm and 35–60% identity (average 48.9%) with gymnosperm NDC80 protein sequences (Table S2). Sequence identity between the peptide immunogen and NDC80 in *R. pubera* was 60%, and the two sequences shared a stretch of five identical amino acids (RMKRE) (Fig. 1). However, as almost the entire *R. pubera* NDC80 domain was composed of amino acid residues similar or identical to the peptide immunogen (Fig. 1, Table S2), it could not be predicted whether the RMKRE motif represents a putative epitope. Moreover, in comparison with the FFGPVS motif found in KNL1, the RMKRE motif of NDC80 was less conserved, being present in 206 of 343 (60%) angiosperm species and absent in all analyzed gymnosperm species and non-seed plants. On the other hand, 276 of the 343 (80.5%) angiosperm species showed ≥60% sequence identity with the peptide immunogen and possessed at least one stretch of ≥5 amino acid residues identical to the peptide immunogen. Since this level of sequence identity was sufficient for centromere labeling in *R. pubera*, it is possible that the anti-NDC80 antibody could also work in many other angiosperm species.

### Confirmation of the efficacy of antibodies by in-situ immunodetection

We performed immunostaining experiments to (1) test the functionality of the antibodies in species with different phylogenetic distances, (2) determine how the antibodies perform in species with different centromere organization, and (3) analyze how the changes in peptide sequence affect the success of centromere labeling. A diverse selection of 16 plant species from 11 different orders were used (Table 1). These plant species represented six families of eudicots, i.e., Convolvulaceae (*Cuscuta reflexa*), Droseraceae (*Dionaea muscipula* and *Drosera capensis*), Lamiaceae (*Ocimum basilicum*), Fabaceae (*Pisum sativum*), Brassicaceae (*Arabidopsis thaliana* and *Raphanus sativus*), and Nelumbonaceae (*Nelumbo nucifera*, representative basal dicot), and four families of monocots, namely, Juncaceae (*Juncus effusus* and *Luzula nivea*), Cyperaceae (*Rhynchospora pubera*), Poaceae (*Triticum aestivum*), and Melanthiaceae (*Chionographis japonica*), including yet the most phylogenetically distant species, namely, *Nymphaea alba* (Nymphaeaceae, representative basal angiosperm), *Picea abies* (Pinaceae, representative gymnosperm), and *Selaginella moellendorffii* (Selaginellaceae, representative non-seed plant). Among these species, *Pisum sativum* served as a representative of those with metapolycentric centromere organization, while *Luzula nivea*, *Rhynchospora pubera*, and *Chionographis japonica* represented the holocentrics. The centromere organization of *Dionaea muscipula* has been inconsistent in the literature, with the species being identified as both monocentric and holocentric. We therefore decided to include *D. muscipula* in this study to resolve its centromere organization.

**Table 1.** Details of plant species included in this study and the conditions used for chromosome fixation and isolation.

| Species | Fixation temperature | Slide preparation / enzymatic digestion | Source of the plant material |
| --- | --- | --- | --- |
| *Arabidopsis thaliana* | 10°C | Suspension | Ecotype Columbia; the European Arabidopsis Stock Centre (Loughborough, UK) |
| *Chionographis japonica* | on ice | Suspension | Iwasaki-engei, Hokaido, Japan;<br>http://www.iwasaki-engei.co.jp |
| *Cuscuta reflexa* | 10°C | Suspension | Botanic Gardens of the Rhenish Friedrich, Wilhelm University, Bonn, Germany |
| *Dionaea muscipula* | 10°C | Suspension | Mgr. Markéta Aubrechtová<br>(Borek 37367, Czech Republic) |
| *Drosera capensis* | 10°C | Squash /<br>60 min - 37°C | FYTO REIDL – MASOŽRAVÉ ROSTLINY<br>(České Budějovice 37010, Czech Republic) |
| *Juncus effusus* | 10°C | Squash /<br>30 min - 27,4°C | Var. spiralis; commercially obtained,<br>https://www.dingers.de/ |
| *Luzula nivea* | on ice | Squash /<br>60 min - 27,4°C | https://www.dingers.de/ |
| *Nelumbo nucifera* | 4°C | Suspension | https://www.levnasemena.cz |
| *Nymphaea alba* | 4°C | Suspension | https://www.bonsai-shop.cz |
| *Ocimum basilicum* | 4°C | Suspension | NOHEL GARDEN a.s.<br>(Dobříš 26301, Czech Republic) |
| *Picea abies* | 4°C | Suspension | Collected from a spruce stand in Želízkův Mlýn, Strážkovice, Czech Republic |
| *Pisum sativum* | 4°C | Suspension | Cv. Terno; Selgen, Stupice, Czech Republic |
| *Raphanus sativus* | 10°C | Suspension | Cv. Lada; MORAVOSEED CZ,<br>Mušlov, Mikulov, Czech Republic |
| *Rhynchospora pubera* | on ice | Squash /<br>50 min - 37°C | Collected in Recife-PE, Brazil. |
| *Selaginella moellendorffii* | 4°C | Suspension | Dr. Iva Mozgová (Biology Centre CAS, České Budějovice, Czech Republic) |
| *Triticum aestivum* | 4°C | Suspension | CZ-BIO-001- PRO-BIO, obch. Spol. s.r.o.,<br>Lipová 40, Staré Mesto, Czech Republic |

The anti-KNL1 antibody exhibited remarkable efficacy, enabling the identification of centromeres in all examined species (Fig. 2) with the exception of *Selaginella moellendorffii*, the representative of non-seed plants (Fig. 2p). This antibody successfully labeled centromeres in many species with monocentric organization, including *Cuscuta reflexa*, *Drosera capensis*, *Ocimum basilicum, Arabidopsis thaliana*, *Raphanus sativus, Nelumbo nucifera*, *Juncus effusus*, *Triticum aestivum, Nymphaea alba*, and *Picea abies,* and produced highly intense and well-defined signals in metapolycentrics (*Pisum sativum*) and holocentrics (*Luzula nivea, Rhynchospora pubera*, and *Chionographis japonica*). In *Dionaea muscipula,* the discrete signals of the anti-KNL1 antibody both on chromosomes and in nuclei indicated a monocentric type of centromere organization (Fig. 2c).

**Fig. 2.**
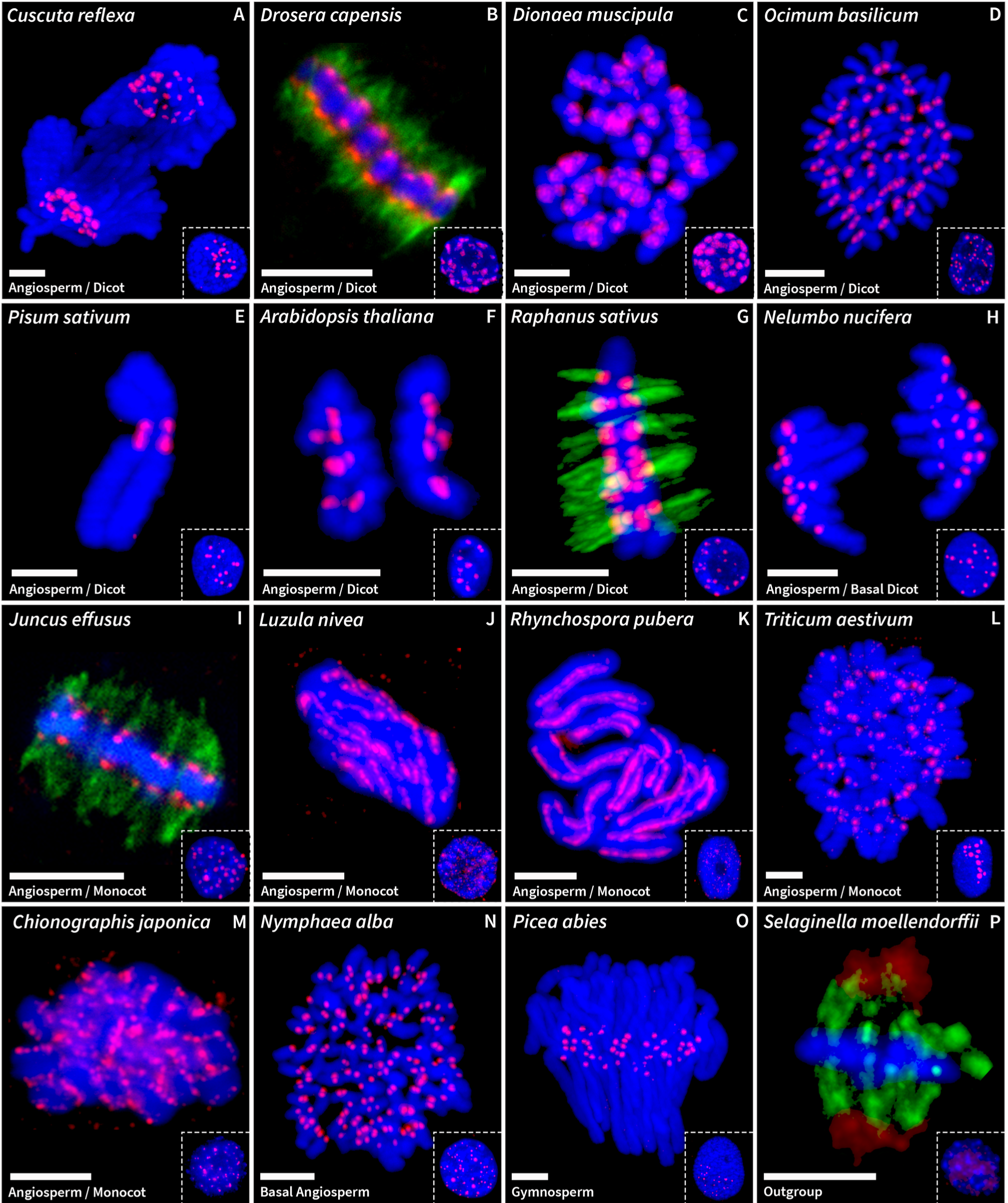
Immunostaining of KNL1 (red) in the chromosomes and interphase nuclei of 16 plant species examined in this study. (A) *Cuscuta reflexa*; (B) *Drosera capensis*; (C) *Dionaea muscipula*; (D) *Ocimum basilicum*; (E) *Pisum sativum*; (F) *Arabidopsis thaliana*; (G) *Raphanus sativus*; (H) *Nelumbo nucifera*; (I) *Juncus effusus*; (J) *Luzula nivea*; (K) *Rhynchospora pubera*; (L) *Triticum aestivum*; (M) *Chionographis japonica*; (N) *Nymphaea alba*; (O) *Picea abies*; (P) *Selaginella moellendorffii*. Green signals in B, G, I and P show immunostaining of α-tubulin. Chromosomes were stained with DAPI (blue). Scale bars, 5 μm.

The anti-NDC80 antibody proved effective for the detection of centromeres in 9 of the 16 species (Fig. 3) but was not effective in *Dionaea muscipula, Ocimum basilicum*, *Arabidopsis thaliana*, *Chionographis japonica, Nymphaea alba, Picea abies*, and *Selaginella moellendorffii*. Although its efficiency was lower compared with anti-KNL1 antibody, the anti-NDC80 antibody labeled centromeres with different types of organization in a wide range of plant species. Notably, the anti-NDC80 antibody signal was more difficult to detect in *Triticum aestivum* (Fig. 3l), particularly on chromosomes that were either not released from cells or covered with cell debris.

**Fig. 3.**
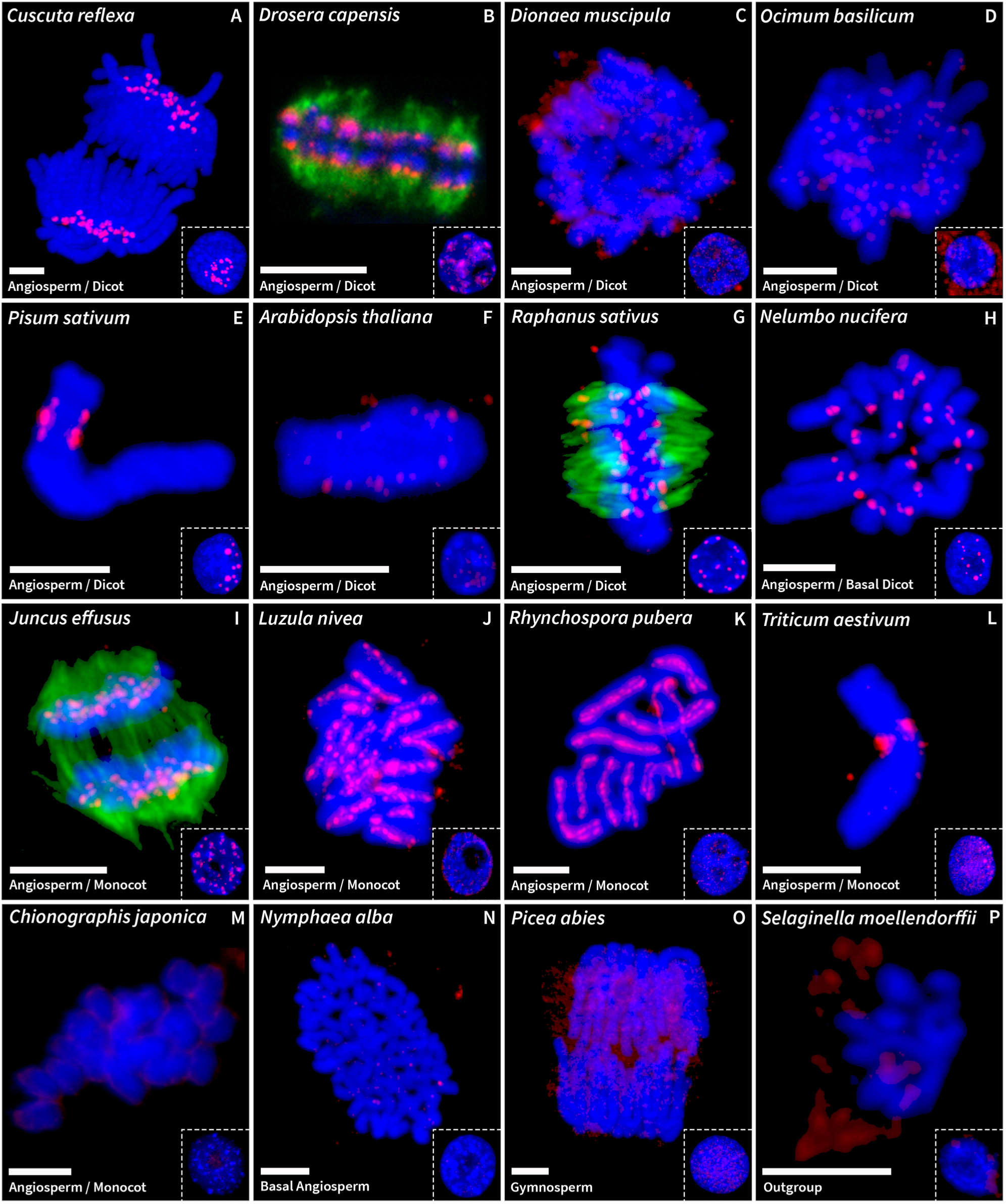
Immunostaining of NDC80 (red) in the chromosomes and interphase nuclei of 16 plant species examined in this study. (A) *Cuscuta reflexa*; (B) *Drosera capensis*; (C) *Dionaea muscipula*; (D) *Ocimum basilicum*; (E) *Pisum sativum*; (F) *Arabidopsis thaliana*; (G) *Raphanus sativus*; (H) *Nelumbo nucifera*; (I) *Juncus effusus*; (J) *Luzula nivea*; (K) *Rhynchospora pubera*; (L) *Triticum aestivum*; (M) *Chionographis japonica*; (N) *Nymphaea alba*; (O) *Picea abies*; (P) *Selaginella moellendorffii*. Green signals in B, G, and I show immunostaining of α-tubulin. Chromosomes were stained with DAPI (blue). Scale bars, 5 μm.

It is worth noting that in species where antibodies labeled centromeres in metaphase chromosomes, both antibodies showed consistent labeling of centromeric chromatin throughout the cell cycle. This resulted in well-defined signals even during interphase (Fig. 2 and Fig. 3), except in the case of holocentric species *Luzula nivea* and *Rhynchospora pubera*, in which the centromeres dissociated into individual units and the signals were scattered whenever the chromatin was not condensed. Another exception was the monocentric species *Picea abies*, in which only ∼25% of the nuclei (n = 100) showed centromeric signals.

Importantly, we noted that the chromosome preparation technique played a crucial role in achieving clear signals during immunostaining. In the case of *Ocimum basilicum* and *Nelumbo nucifera*, centromeric signals could not be detected using either antibody when chromosomes were prepared using the squash technique. However, a reliable detection of centromeres was achieved with the anti-KNL1 antibody in *O. basilicum* and with both antibodies in *N. nucifera* when chromosomes were isolated into a suspension. Both chromosome preparation techniques were also tested in *Cuscuta reflexa, Dionaea muscipula, Pisum sativum, Raphanus sativus, Triticum aestivum*, and *Picea abies*. Although squashed chromosomes showed centromeric signals in some species, the results were significantly improved and more reproducible with the chromosome suspension method.

## Discussion

In this study, we showed that immunostaining centromeres by targeting conserved domains in the outer kinetochore proteins KNL1 and NDC80 provides the most versatile means, to date, for identifying functional centromere domains in seed plants. The anti-KNL1 antibody proved to be particularly versatile and efficient. By combining immunostaining results with amino acid sequence alignments, we found that this antibody could reliably recognize centromeres even in the gymnosperm *Picea abies*, in which the target domain, whose sequence is considerably different from that of the peptide immunogen, shares a short motif of six identical amino acids (FFGPVS). This indicates that the FFGPVS motif is a possible epitope. The labeling of centromeres in *Pisum sativum* and *Triticum aestivum*, each of which has one substitution in the motif, albeit at a different position, suggests that sequence variation in the motif does not necessarily prevent the antibody from binding to the target domain (Figs. 1 and 2). Considering that the FFGPVS motif was found in 83% of angiosperm species and all gymnosperm species included in this study, and that the anti-KNL1 antibody also functioned in species harboring variation in the motif sequence, it is very likely that the anti-KNL1 antibody developed in this study is capable of labeling centromeres in the vast majority of seed plants.

Although the anti-NDC80 antibody could label centromeres in species from evolutionarily diverse lineages, such as monocots and dicots, its versatility was lower than that of the anti-KNL1 antibody. This may be due to the combined effect of two factors: 1) sequence divergence of the epitope(s) in the target domain, and 2) high proportion of amino acid residues that can be chemically modified by formaldehyde in the fixative, so that they are no longer recognized by the antibody (Metz et al. 2004, Fig. S2). The latter seems likely to have been the case in *Ocimum basilicum*, where the anti-NDC80 antibody failed to label centromeres, although the similarity of the target domain to the peptide immunogen was higher than in *Pisum sativum*, for example, where the antibody functioned very well. By contrast, formaldehyde-reactive amino acid residues were less abundant in the target domain of KNL1 and were particularly absent in its FFGPVS motif (Fig. S2). The higher sensitivity of the NDC80 target domain to formaldehyde therefore places higher demands on the optimization of the fixation conditions.

Immuno-detection of kinetochore proteins is the most accurate approach for the identification of functional centromeres. This is because kinetochores form specifically at functional centromere domains, whereas other markers such as various types of histone phosphorylation tend to occur at pericentromeres (Zhang et al. 2014). Since H2AT120ph has been detected in the chromosomes of many plant species, the antibody directed against this marker was considered the most universal for centromere labeling (Demidov et al. 2014; Jankowska et al. 2015; Wanner et al. 2015; Báez et al. 2019). However, the anti-H2AT120ph antibody is not an ideal replacement for the anti-CENH3 antibody for several reasons: 1) H2AT120ph is not only enriched in centromeres but also in pericentromeric regions (Wanner et al. 2015; Neumann et al. 2016; Báez et al. 2019); 2) H2AT120ph has also been detected in small amounts on chromosome arms (Demidov et al. 2014; Báez et al. 2019); 3) H2AT120ph shows a non-centromeric distribution during meiosis in *Rhynchospora pubera* (Cabral et al. 2014); and 4) when detected together with CENH3, H2AT120ph is seen more toward the inner region of the centromere, whereas CENH3 is more at the peripheral regions (Demidov et al. 2014; Neumann et al. 2016), with the two usually localized to different nucleosomes.

The signals produced by anti-KNL1 and anti-NDC80 antibodies in both mitotic chromosomes and interphase nuclei indicated that the outer kinetochore proteins occur at centromeres throughout the cell cycle. This feature provides an extra advantage to the study of centromeres in plants. During interphase, monocentromeres in plants typically manifest as distinct, dense bodies referred to as chromocenters (Fransz et al. 2006; Lermontova et al. 2015), a characteristic mirrored by the discrete signals of both antibodies in the nuclei of monocentric species. By contrast, holocentromeres, such as those observed in *Luzula nivea* (Figs. 2j and 3j; Nagaki and Murata 2005) and *Rhynchospora pubera* (Figs. 2k and 3k; Marques et al. 2015), dissociate into individual units, resulting in scattered signals. In *Chionographis japonica*, a functionally holocentric species with 7–11 evenly spaced megabase-sized centromere units per chromosome, centromeres remain organized as discrete loci, outnumbering the chromosomes (Figs. 2m and 3m; Kuo et al. 2023). On the other hand, centromere domains of metapolycentric chromosomes in *Pisum sativum*, a functionally monocentric species with multiple centromere domains per chromosome, merged during interphase into clusters equal or fewer than the number of chromosomes (Figs. 2e and 3e; Neumann et al. 2012; Macas et al. 2023). Therefore, the visualization of kinetochore proteins during interphase proves beneficial for determining the organization and dynamics of centromeres throughout the cell cycle.

Among the species subjected to immunolocalization experiments in this study, we chose *Dionaea muscipula* to scrutinize centromere organization, because of the existing inconsistencies regarding its centromere organization in the literature. While Hoshi and Kondo (1998) assumed that *D. muscipula* is monocentric based on the observation of chromosome morphology and CMA/DAPI banding patterns, Kolodin et al. (2018) determined its centromere type using flow cytometry analysis of nuclei from irradiated plants and assumed holocentricity. Our analysis of KNL1 in *D. muscipula* indicated that the species is monocentric, since the signals occurred at discrete loci both on chromosomes and in nuclei (Fig. 2c).

We found that the detection of kinetochore proteins was significantly enhanced when chromosomes were prepared using the suspension method instead of the squash technique. We believe that this discrepancy was not caused by the unavailability of epitopes due to cross-linking, since the fixation conditions were the same for both methods, but by the enzymatic digestion required to remove the cell wall in the squash method. If the material is not sufficiently digested, it may be difficult for the antibody to penetrate the remaining cell wall and cytoplasm. Conversely, if the material is over-digested, the integrity of cells may be compromised (Brown and Lemmon 1995), which may affect the structure of kinetochores. Therefore, we believe that the detection of kinetochore proteins can be improved by determining the optimal digestion time. However, based on our observations, we recommend isolation of chromosomes into a suspension as the more appropriate method for preparing chromosomes when immunostaining kinetochore proteins.

Although the polyclonal nature of the antibodies developed in this study does not allow their exact replication, our results showed that the domains used as peptide immunogens are highly conserved in sequence and are immunogenic in seed plants, which suggests that future attempts at developing antibodies against these proteins will be successful. In addition, the FFGPVS motif in KNL1 provides a unique opportunity for developing a highly versatile monoclonal antibody against this protein. While CENH3 will likely continue to be the better centromere-specific target protein for some techniques, such as chromatin immunoprecipitation, our results demonstrate that KNL1 and NDC80 are better suited for immunolabeling centromeres, because antibodies against these proteins offer incomparably greater versatility across different plant species, a finding that is particularly convenient for studying the organization and function of the centromere in non-model species.

## Material and methods

### Plant material

The origin and details of plant species used in this study are described in Table 1.

### Generation of KNL1- and NDC80-specific antibodies

Antibodies against KNL1 and NDC80 proteins were produced using peptide immunogens EDHFFGPVSPSFIRPGRLSDC and CQGINARDAERMKRELQALEG, respectively. The peptides synthesis, immunization of rabbits and peptide affinity purification of antisera were performed by GenScript (KNL1; Piscataway, NJ, USA) and Biomatik (NDC80; Cambridge, ON, Canada). Additional details of the sequences used to design the peptide immunogens can be found in our previous study (Neumann et al. 2023).

### Sequence analyses

Homologs of KNL1 and NDC80 proteins were identified through blastp and tblastn searches (Altschul et al. 1997) using the amino acid sequences of these two proteins identified previously in *Cuscuta* spp., *Ipomoea* spp., and *Arabidopsis thaliana* (Neumann et al. 2023) as primary queries. Accession numbers and source databases of all sequences are provided in Tables S1 and S2. Sequence alignments were performed using MUSCLE (Edgar 2004) and edited in SEAVIEW (Galtier et al. 1996). Sequence logos were generated using WebLogo (Crooks et al. 2004).

### Chromosome preparation and immunostaining

To prepare the nuclei and chromosomes for immunostaining, different biological materials were utilized depending on the plant species: shoot tips for *Cuscuta reflexa*, root tips and young leaves for *Arabidopsis thaliana*, and root tips for the remaining species. Pre-treatment was performed only for *Chionographis japonica* (ice-cold water at 4°C overnight) and *Pisum sativum* (1.25 mM hydroxyurea and 15 µM oryzalin, according to Neumann et al. 2002). The following information on slide preparation and immunostaining applies to all species, with the exception of *Luzula nivea*, *Rhynchospora pubera*, and *Chionographis japonica*, for which the methodology described in Marques et al. (2015), Marques et al. (2016) and Kuo et al. (2023), respectively, should be consulted. Chromosomes were fixed in 3% formaldehyde diluted in Tris-fix buffer (10 mM Tris, 10 mM Na_2_EDTA, 100 mM NaCl [pH 7.5]), with the first 5 min under vacuum, and then washed with Tris buffer on ice for 30 min. The fixation temperature, chromosome preparation method (suspension or squash), and enzymatic digestion conditions varied with the plant species (Table 1). When using the squash technique, the biological material was digested using 2% cellulase ONOZUKA R10 (SERVA Electrophoresis, Heidelberg, Germany) and 2% pectinase (MP Biomedicals, Santa Ana, CA, USA) for varying durations (Table 1), and the squashes were performed in 1× phosphate-buffered saline (PBS). When preparing chromosomes and nuclei using the suspension technique, the tissue was ground in 1 ml of cold LB01 (15 mM Tris, 2 mM Na 2 EDTA, 80 mM KCl, 20 mM NaCl, 0.5 mM spermine, 15 mM mercaptoethanol, 0.1% Triton X-100 [pH 7.5]) using a mechanical homogenizer (Ultra-turrax T8, IKA Z404519), and the resulting suspension was filtered through a 48-μm nylon mesh and deposited onto slides using a centrifuge with cytospin chambers (Hettich). Slides were washed once with 1× PBS for 5 min, and then incubated at room temperature (RT) in 1× PBS with 0.5% Triton (pH 7.4) for 30 min before immunostaining to improve permeabilization. This step was followed by two washes in 1× PBS for 5 min at RT and one wash in 1× PBS with Tween20 (1× PBS, 0.1% Tween20 [pH 7.4]) for 5 min at RT. To conduct immunostaining, slides were incubated with the primary antibody diluted in 1× PBS with Tween20 overnight at 4°C. The dilution ratios were 1:1000 and 1:100 for antibodies against the kinetochore proteins and α-tubulin, respectively (Sigma-Aldrich, St. Louis, MO; catalog number T6199). On the next day, the slides were subjected to two 5-min washes with 1× PBS at RT and one 5-min wash with 1× PBS with Tween20 at RT. Subsequently, the slides were incubated with the secondary antibody in 1× PBS with Tween20 at RT for 1 h, and then washed twice for 5 min in 1× PBS at RT. Primary rabbit and mouse antibodies were detected with goat anti-rabbit Rhodamine Red X (1:500 dilution; Jackson ImmunoResearch, Suffolk, UK; catalog number: 111-295-144) and goat anti-mouse Alexa Fluor 488 (1:500 dilution; Jackson ImmunoResearch; catalog number: 115-545-166), respectively. The slides were post-fixed in 4% formaldehyde diluted in 1× PBS for 10 min at RT, counterstained with 4′,6-diamidino-2-phenylindole (DAPI), and mounted in Vectashield mounting medium (Vector Laboratories, Burlingame, CA). All pictures were taken with the conventional wide-field fluorescence microscope Zeiss AxioImager.Z2 microscope equipped with an AxioCam 506 mono-color camera and with an Apotome2.0 device, except for the images of *Chionographis japonica,* taken by an epifluorescence microscope BX61 (Olympus Europa SE &Co. KG, Germany) equipped with an Orca ER CCD camera (Hamamatsu, Japan).

## Acknowledgements

This research was financially supported by grants from the Czech Science Foundation (20-25440S) and the Czech Academy of Sciences (RVO:60077344). AM is financially supported by the Max Planck Society and Deutsche Forschungsgemeinschaft (grant number MA 9363/3-1). YMS was financially supported by a PROBRAL - Coordenação de Aperfeiçoamento de Pessoal de Nível Superior / Deutscher Akademischer Austauschdienst grant (number 495995/2020-00 and 88881.144086/2017-01). Computational resources and data storage facilities were provided by the ELIXIR-CZ Research Infrastructure Project (LM2018131). We thank J. Látalová and V. Tetourová for technical assistance.

## Authors contribution

LO, PN and JM conceived the study and designed the experiments. LO, YMS and YTK performed the cytogenetics experiments and conventional fluorescence microscopy. PN analyzed the sequenced data. AM provided *Luzula nivea* sequencing data. LO and PN wrote the manuscript with input from YMS, YTK, AM, VS and JM. All authors read and approved the final manuscript.

## Declarations

### Ethical approval

Not applicable.

### Consent to participate

Not applicable.

### Consented for publication

Not applicable.

---

**Fig. S1.**
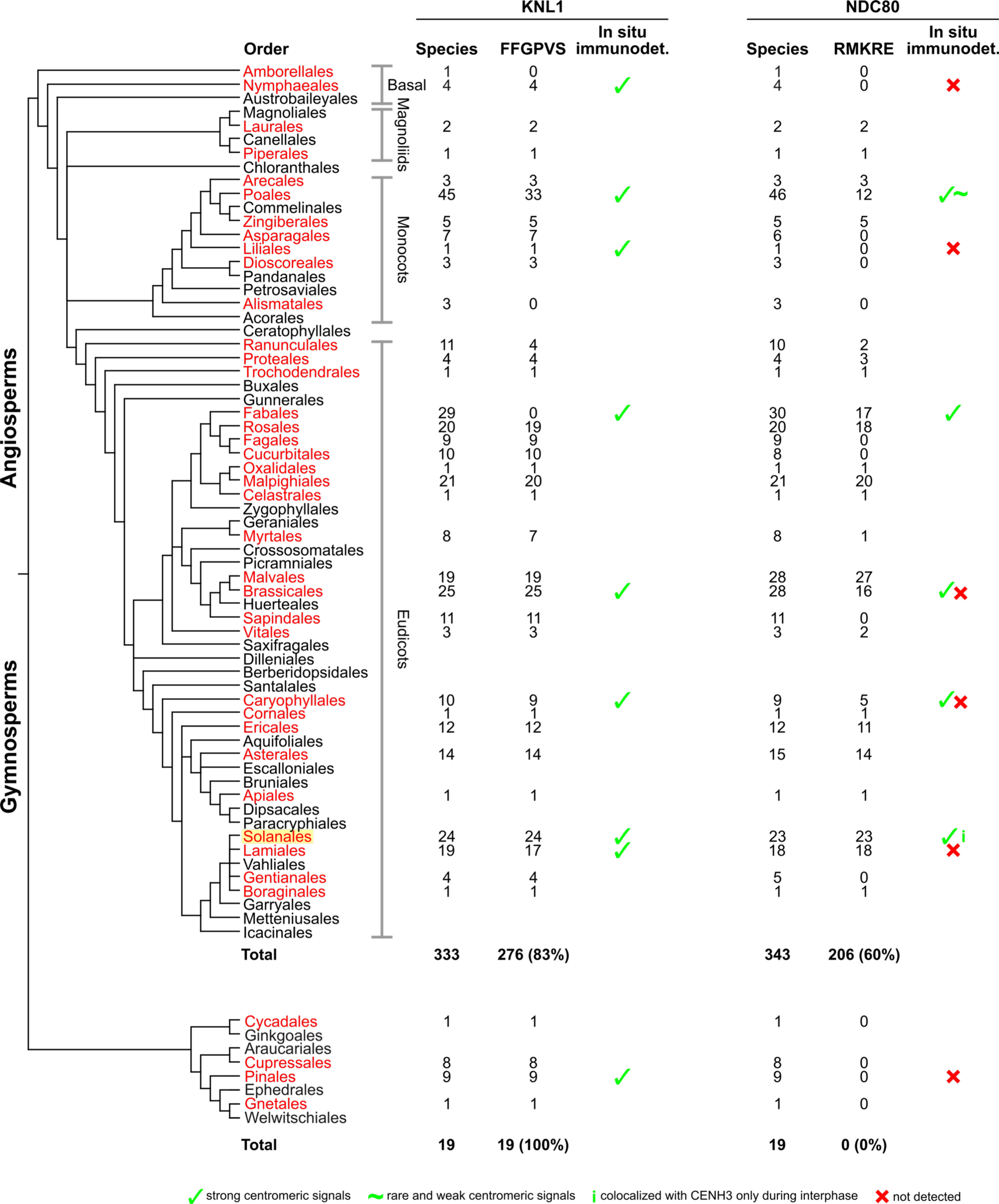
Phylogeny of seed plants. Orders represented in KNL1 and NDC80 sequence datasets are marked in red. *Cuscuta* species, which were the primary targets of antibodies, belong to the order Solanales (highlighted in yellow). Columns on the right side show the number of analyzed species in each order and the number of species possessing the conserved motifs in KNL1 (FFGPVS) and NDC80 (RMKRE). The phylogenetic tree was drawn according to The Angiosperm Phylogeny Group 2016 (Yang et al. 2022).

**Fig. S2.**
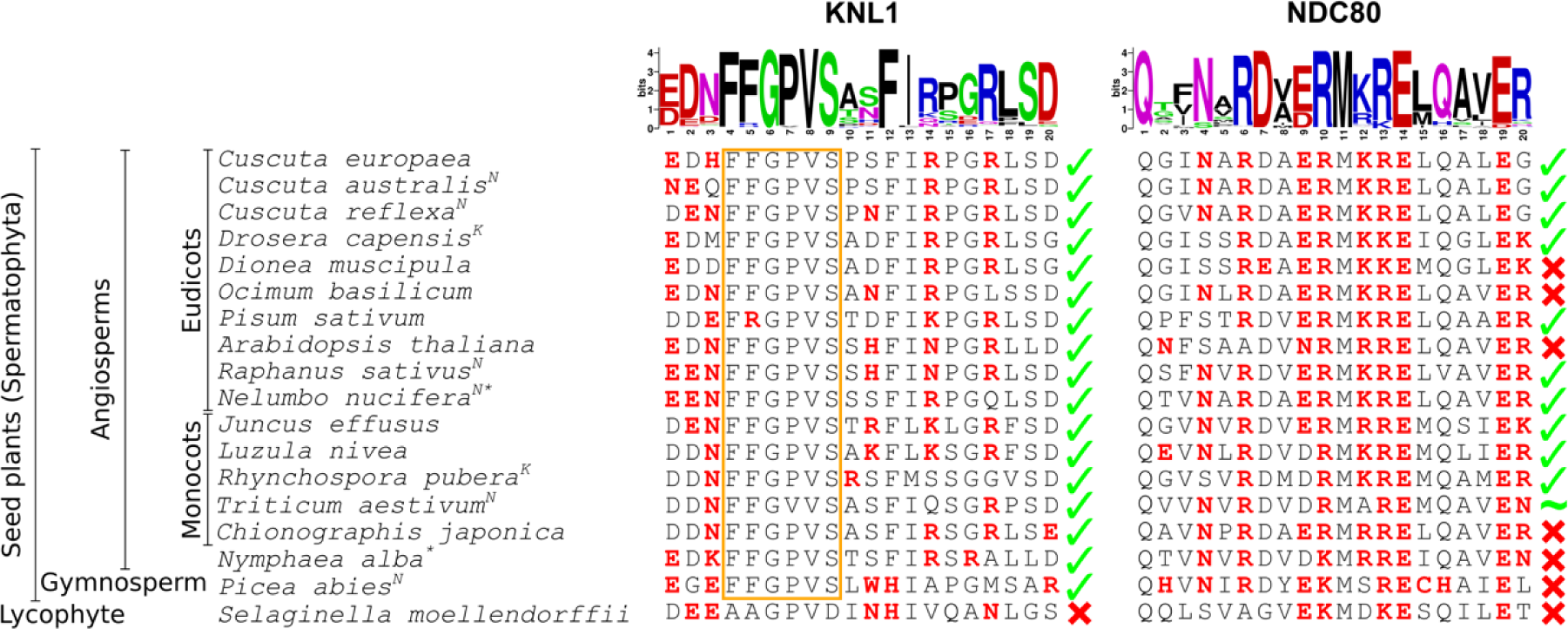
Formaldehyde-reactive amino acid residues in the target domains of KNL1 and NDC80. This figure was modified from Fig. 1 to show amino acid residues (red) that can react with formaldehyde directly or with formaldehyde reaction adducts (Metz et al. 2004). Note that none of the amino acid residues in the conserved motif of KNL1 (FFGPVS) were predicted to react with formaldehyde, which suggests that the putative recognition of the motif by the anti-KNL1 antibody is insensitive to the stringency of the fixation conditions. This is in contrast to NDC80, which contains numerous formaldehyde-reactive amino acid residues along the entire target domain and therefore requires fine-tuning of the chromosome fixation conditions to allow sufficient stability of the chromosomes without excessive modification of the target domain.

